# P-GRe : an efficient pipeline to maximised pseudogene prediction in plants/eucaryotes

**DOI:** 10.1101/2023.12.04.569967

**Authors:** Sébastien Cabanac, Christophe Dunand, Catherine Mathé

## Abstract

Formerly considered as part of “junk DNA”, pseudogenes are nowadays known for their role in the post-transcriptional regulation of functional genes. In addition, their identification allows a better understanding of gene evolution in the frame of multigenic families. Despite this, there is, to our knowledge, no fully automatic user-friendly software allowing the annotation of pseudogenes on a whole genome. Here, we present Pseudo-Gene Retriever (P-GRe), a fully automated pseudogene prediction software requiring only a genome sequence and its corresponding GFF annotation file. P-GRe detects the sequences of the pseudogenes on a whole genome and returns to the user all their genomic sequences and their pseudo-coding sequences. The ability of P-GRe to finely reconstruct the structure of pseudogenes also allow to obtain a set of proteins virtually encoded by the predicted pseudogenes. We show here that in 70% of the cases, virtual proteins constructed by P-GRe from *Arabidopsis thaliana* proteome and genome aligned better to their parent protein than their annotated counterpart.

## INTRODUCTION

Pseudogenes are genomic sequences with homology to functional genes but that harbor deleterious mutations, such as loss of the start codon, loss of coding sequence part, gain of stop or frame-shifts. No longer coding for a functional protein, pseudogenes are rarely transcribed and are often described as having no function. It is known that part of the pseudogenes are transcribed, this part representing for example 15% of all the pseudogenes in mice (Sisu *et al*., 2020). It has also been shown that these transcripts, originating from pseudogenes, could form duplexes with the mRNAs of homologous functional genes and thus participate in post-transcriptional regulation through the RNAi pathway (Tam *et al*., 2008; Watanabe *et al*., 2008; Guo *et al*., 2009). Moreover, the exhaustive prediction of pseudogenes could allow a better understanding of dynamic of gene evolution in multigenic families often subjected to duplication and pseudogenization events.

Most of the current pseudogene prediction software relies on the homology between the known protein sequences of an organism and the sequences of the pseudogenes to predict the positions of these pseudogenes on the genome. Briefly, one or more local alignments of each protein sequence are performed in order to find an approximate position of the pseudogenes. The protein sequence with the highest similarity with the sequence found for each local alignment is defined as being encoded by the parent gene, as it is assumed that many pseudogenes are derived from the duplication of functional genes. A finer alignment is then carried out between the hits obtained at the end of the first local alignment and the associated parent sequences. Shiu’s pipeline (Zou *et al*., 2009) and PseudoPipe (Zhang *et al*., 2006), two software commonly used for pseudogene prediction, compares protein sequences to DNA sequence database in both DNA orientations using the tfasty local alignment algorithm (Pearson & Lipman, 1988).

If a large number of pseudogene prediction software exist, many are specific to a type of organism, such as Pseudofinder (Syberg-Olsen *et al*., 2022) and Psi-Phi (Lerat & Ochman, 2004) which are dedicated to prokaryotes. Others target specific types of pseudogenes, such as PPFINDER (van Baren & Brent, 2006) and PΨFinder (Abrahamsson *et al*., 2022) which are made to predict pseudogenes originating from transcript retrotransposition events. Software able to work on any type of organisms and pseudogenes are rarer, and often produce very different outputs: some can approximate a protein or a peptide virtually encoded by the pseudogenes, while others simply return the sequence of the pseudogenes without worrying about their pseudo-coding structure. Most generate their own results file in TSV format and surprisingly, to our knowledge, none of the software returns its results in the GFF or GTF reference formats mainly used for structural genome annotation. Last, these software are often not very user-friendly, requiring a tedious preparation of the data and the organization of the working directories according to precise instructions.

Pseudo-Gene Retriever (P-GRe) allows the prediction of the positions of pseudogenes on a whole genome as well as the precise reconstruction of their structures in pseudo-coding exon. P-GRe only requires the genome sequence and the corresponding structural annotation in GFF format. It is therefore optional to provide all the protein sequences, as P-GRe can generate them from genome and annotation files. P-GRe returns the set of genomic sequences of the pseudogenes, and also the set of pseudo-coding sequences. The positions of pseudogenes and their features on the genome are returned in GFF format, including pseudo-exons, start and stop codons as well as frame-shifts. Precise reconstruction of the pseudogenes’ structure and detection of frame-shift events also enable P-GRe to return the list of proteins and peptides virtually encoded by the predicted pseudogenes. The P-GRe process integrates a set of new approaches, including the detection of frame-shifts by “chimera generation”, as described in the Materials & Methods section, and the replacement of the local alignments, during the step of fine alignment, by global alignments. These alignments are corrected and refined by an algorithm inspired by Lindley’s process (Lindley, 1952) and by a search for canonical splice sites, both described in the Materials & Methods section.

## MATERIALS & METHODS

### P-GRe

Like most pseudogene prediction software, P-GRe proceeds in two steps, the first consisting in finding the approximate position of the pseudogenes and the second in refining their structure (Figure 1). For the first step, P-GRe uses the GFFread software (Pertea & Pertea, 2020) to generate the set of proteins from the genome and the annotation file provided by the user. These same files are used for hard masking annotated gene sequences on the genome using maskfasta from the BEDTools suite (Quinlan & Hall, 2010). The generated proteins are then locally aligned to the masked genome using the tblastn algorithm (Altschul *et al*., 1990), with the options -seg ‘yes’, -soft_masking True, -db_soft_mask dust, - outfmt 6, -evalue 0.01, -word_size 3, -gapextend 2 and -max_intron_length 10000. The local alignments obtained by tblastn are then selected on their percentage of identity. The latter must be greater than a threshold to be selected, this threshold being dependent on the alignment length and calculated by the following formula:

**Figure 1.**
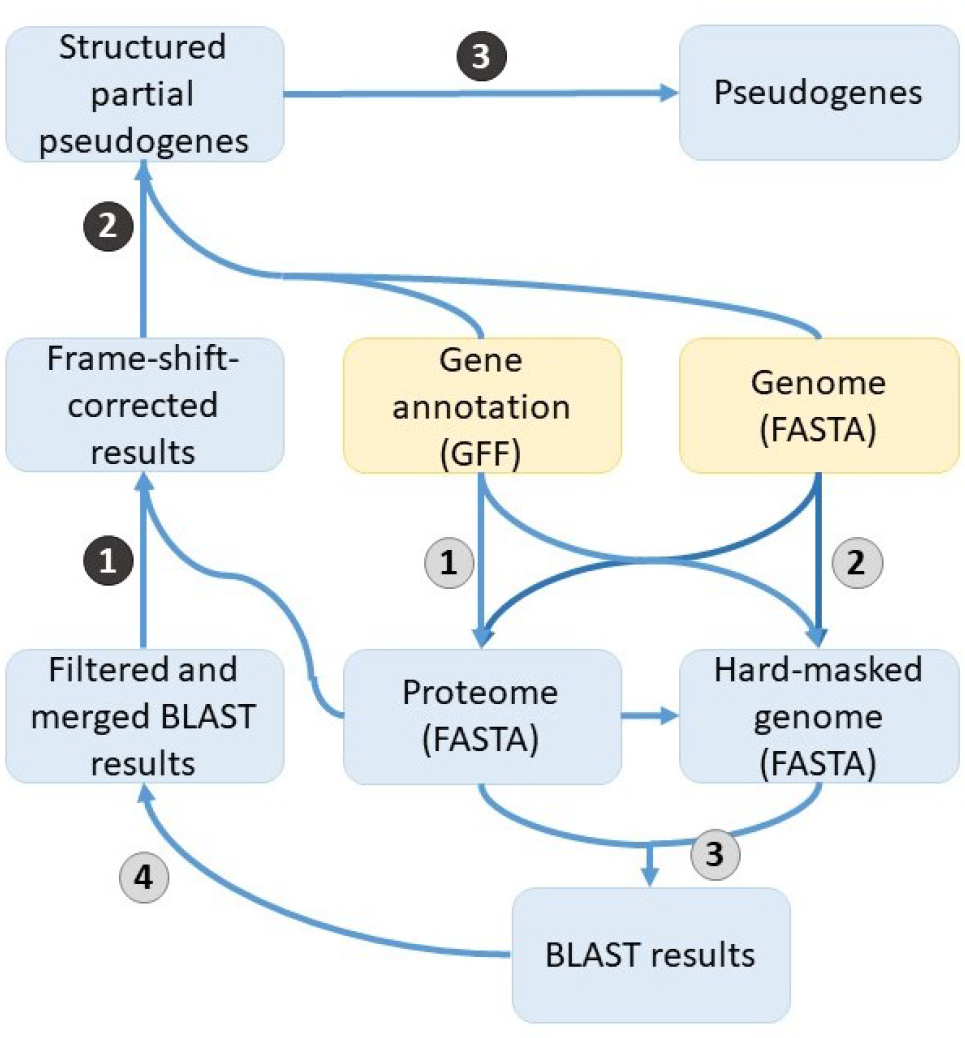
Diagram representing the operation of P-GRe. Mandatory inputs are colored in yellow. The first step of P-GRe, *i*.*e*. the detection of pseudogenes, is made up of four sub-steps numbered in gray circles: 1. Generation of the proteome, 2. Hard-masking of genes on the genome. 3. Local alignments of proteins on the masked genome and 4. Filtering and merging of alignment results. The second step of P-GRe, *i*.*e*. the reconstruction of the pseudogenes, consists of three sub-steps numbered on black circles: 1. Detection of the frame-shifts, 2. Determination of the structure of the pseudogenes and 3. Reconstruction of the terminal parts.

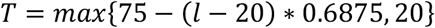

where *T* is the identity threshold and *l* is the length of the alignment. If several local alignments have been obtained with the same protein and are separated by less than 10 kb, all these local alignments are selected. In the end, all non-selected results are filtered out. Remaining overlapping alignments obtained from different proteins are filtered in order to keep only the longest alignment.

The second step, *i*.*e*. the prediction of the structure of each pseudogene, is itself divided into four stages.

First, frame-shifts are identified from overlapping local alignments obtained by the same protein, where the length of the overlap is not divisible by three. The position of the frame-shift is precisely determined after translation in the different reading frames of the local alignments’ area (Figure 2). All of the peptides obtained, called chimeras, are aligned locally with blastp algorithm with the protein sequence coded by the parent gene of the pseudogene. The chimera that aligns best, *i*.*e*. that achieves the lowest E-value, is used to determine the correct frame and the position of the frame-shift. For this step, the following parameters are set in the blastp algorithm: -evalue 0.01, -word_size 3, -gapextend 2, -max_target_seqs 1.

**Figure 2.**
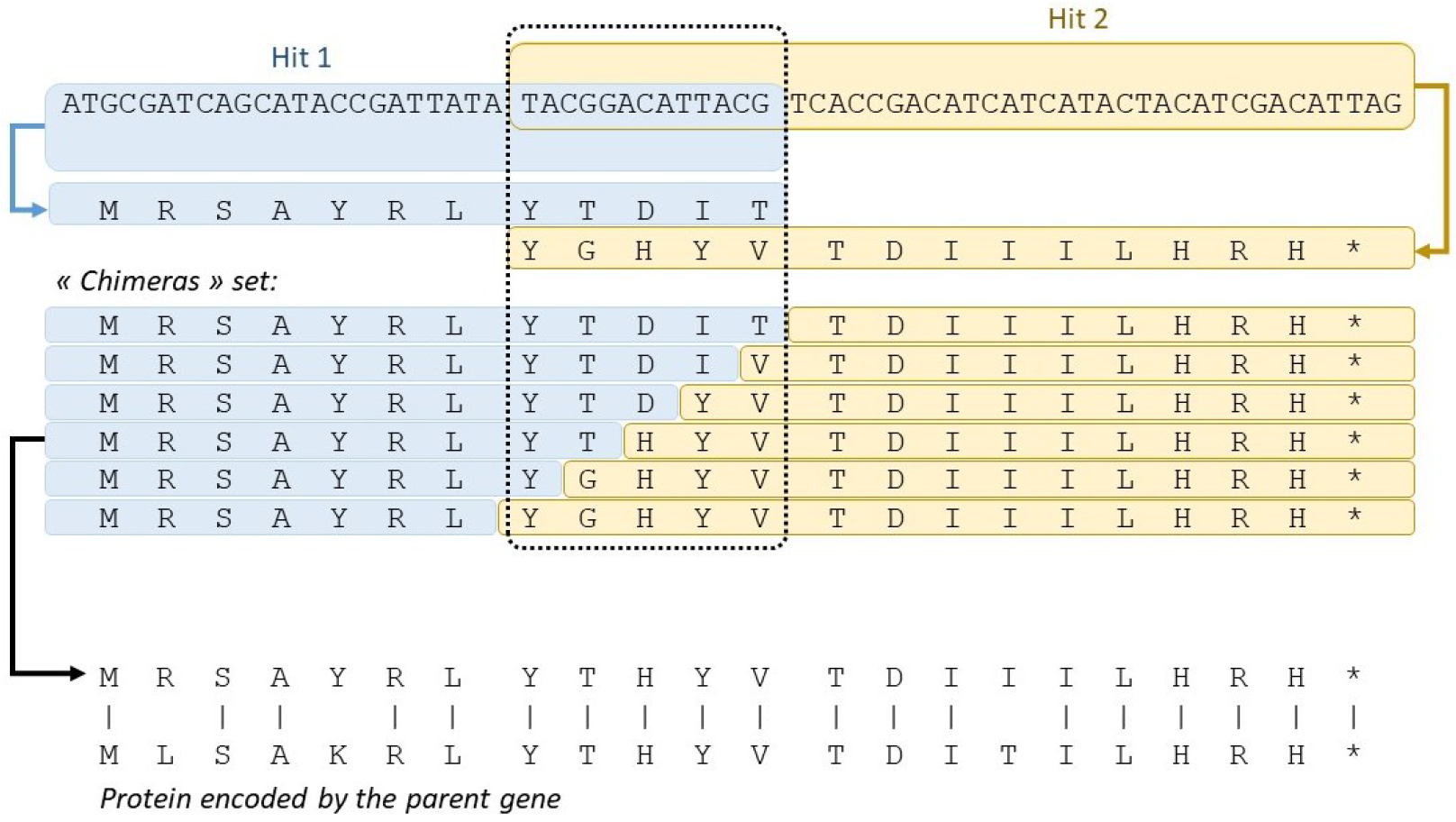
Example of frame-shift detection via “chimera construction”. When two local alignments originating from the same protein overlap, the peptides encoded by the sequences corresponding to these alignments are generated. The two peptides are concatenated, with the amino acids encoded by the overlapping positions of the N-ter side peptide (in blue) being gradually replaced by the overlapping amino acid of the C-ter side peptide (in orange). All the chimeras thus generated are aligned locally on the protein encoded by the parent gene of the pseudogene. The chimera returning the best alignment (the one with the lowest E-value) is then used to determine the position of the frame shift.

Secondly, because local alignments obtained in the first step generally do not cover the entire pseudo-coding exon, a few bases at the ends or the beginning of the pseudo-coding exon may be missing. To overcome this problem, P-GRe extends the positions of the alignments until the next alignment (Figure 3A). The expanded regions are then translated into a single amino acid sequence that is globally aligned with the parent protein sequence. The extended gaps thus generally correspond to the pseudo-introns on the pseudogene. Certain misalignments can falsify this structure, and are corrected by a process inspired by the Lindley process, which is therefore called hereafter “*a priori* Lindley-inspired process” (Figure 3B).

**Figure 3.**
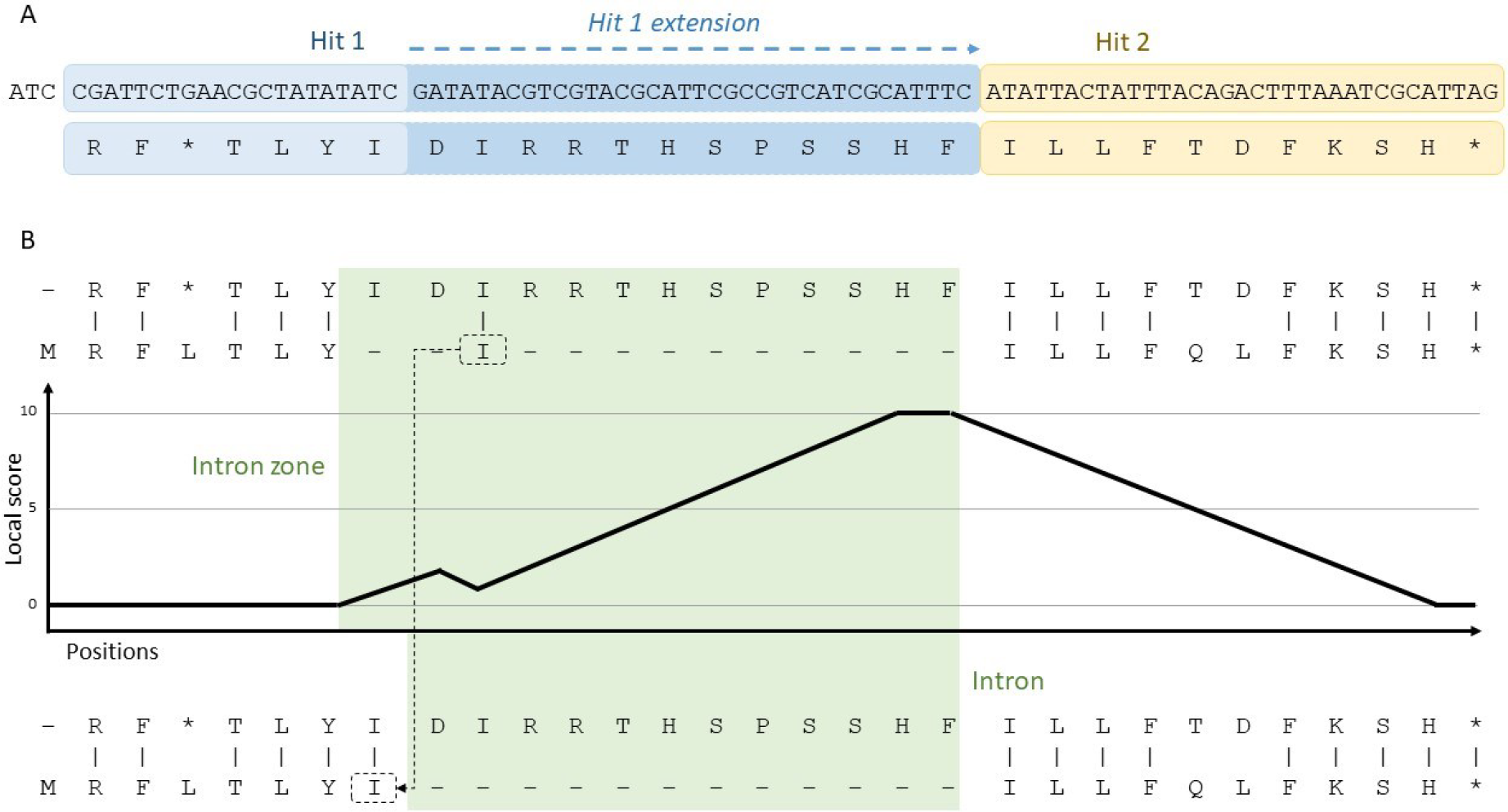
Example of alignment correction performed by P-GRe with a Lindley-inspired process. A. Each local alignment obtained from a protein (hit) is extended to the next hit. B. The resulting peptide is aligned with the protein encoded by the parent gene of the pseudogene. A process inspired by the Lindley process assigns each position a local score, between 0 and 10, equal to the local score of the previous position, +1 if a gap is present on the aligned parent sequence and -1 otherwise. Zones of introns are identified by a sequence of positions with a non-zero score, with the last position having the maximum score. Finally, the positions within the regions of introns to which amino acids are aligned are realigned, the amino acids of the protein encoded by the parent gene being reassembled with the amino acids encoded by the same coding sequence.

Briefly, the alignment between the supposed amino acid sequence and the parent protein sequence is scanned from left to right. Each amino acid coded by the pseudogene is assigned a local score equal to the score of the preceding amino acid, +1 if the latter is aligned with a gap, -1 otherwise. The local score of an amino acid can never be negative and never exceed 10. Along the alignment, local score “spikes” can thus be obtained and are used to define zones of introns, *i*.*e*. the longest series of positions with non-zero scores whose first position has a score equal to 1 and last position has the highest score of the series, this score being greater than or equal to 5. Furthermore, thanks to the information present in the GFF file, the *a priori* Lindley-like process is activated at the approximate positions where an intron is expected, *i*.*e*. five amino acids before each pseudo-coding exon change. The local score can only vary when the process is activated, and the process is deactivated when the local score reaches 0 after defining an intron zone, or if no gaps are found within 10 positions after an activation. For this step, the alignments are carried out using the pairwise2 module of the BioPython tool suite if the two sequences to be aligned are less than 300 amino acids (Cock *et al*., 2009), and using stretcher from the EMBOSS tool suite (Rice *et al*., 2000) otherwise to be faster. As large gaps are expected in low numbers, corresponding to introns, the alignments are performed with a strong gap opening penalty (5) and no extension penalty. The BLOSUM62 matrix is used for substitution scoring. Amino acids encoded by the pseudogene aligning with those of the parent protein within an intron zone are then considered as misalignments.

Thanks to the information present in the GFF file, P-GRe can then reattach a misaligned amino acid from the parent protein sequence to the amino acids encoded by the same coding exon, correcting the alignment. Finally, to refine the structure of the pseudogenes, a search for canonical GT/AT splicing sites is carried out at more or less 9 bases from each start and end of introns.

The third stage consists in reconstructing the N-terminal and C-terminal parts of the proteins virtually coded by the pseudogenes. Thanks to the local alignments carried out during the search for the pseudogenes, P-GRe determines for each pseudogene the positions at which a start codon and a stop codon should be found (Figure 4A). Concerning the start codon, P-GRe will search for an ATG codon or a degenerate ATG codon, *i*.*e*. with one substitution allowed. If no start codon is found at the expected position, P-GRe will search for ATG codons from the expected position towards the start of the first pseudo-coding exon. P-GRe also searches for a stop codon from the expected position towards the end of the last pseudo-exon (Figure 4B).

**Figure 4.**
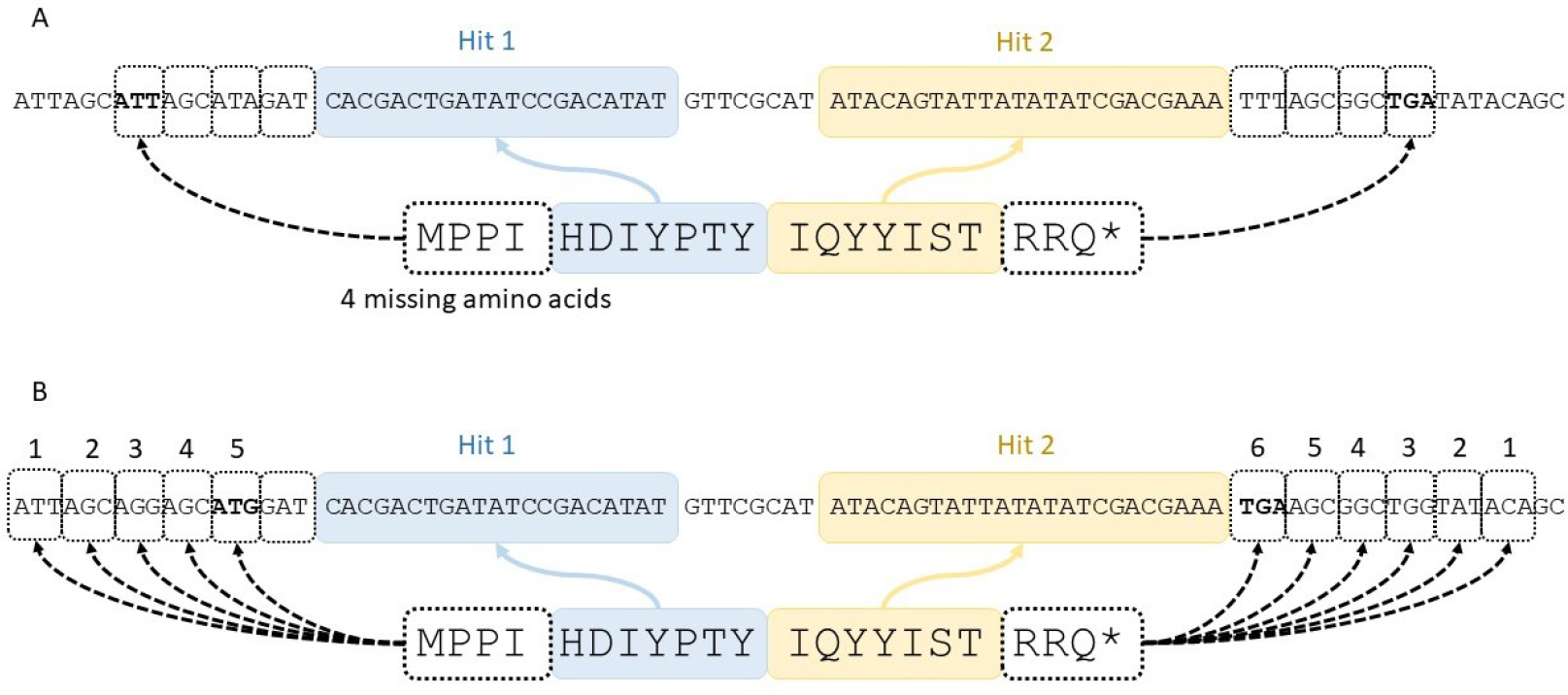
Examples of reconstructions of pseudogene ends by searching for start and stop codons. A. Case where a degenerated start codon is found (ATG codon with a substitution) at the expected position, and where a stop codon is also found at the expected position. B. If no start or degenerate start codon is found, P-GRe will look for a start codon a bit further and gradually look for one, base by base, towards the start of the first exon. For the stop codon, this search is done base by base towards the end of the last exon.

Fourth, once the set of pseudogenes has been predicted, the pseudogenes separated by less than 2.5 kb and with no terminal stop codon are merged. This step makes it possible in particular to reconstitute the pseudogenes whose different exons initially matched to different parent proteins highly similar. This also allows to consider the rare cases of chimeric pseudogenes, *i*.*e*. pseudogenes consisting of fragments of sequences originating from different genes.

### Pseudogenes categorization

The calculations carried out by P-GRe during the different steps are used to categorize the different predicted pseudogenes. Pseudogenes are categorized according to two characteristics: their completeness and their type. Completeness can be noted as “Copy” if the pseudogene appears to be a full copy of the parent gene or “Fragment” if it appears to be only a copy of part of it. This characteristic is determined using a decision tree (Figure 5).

**Figure 5.**
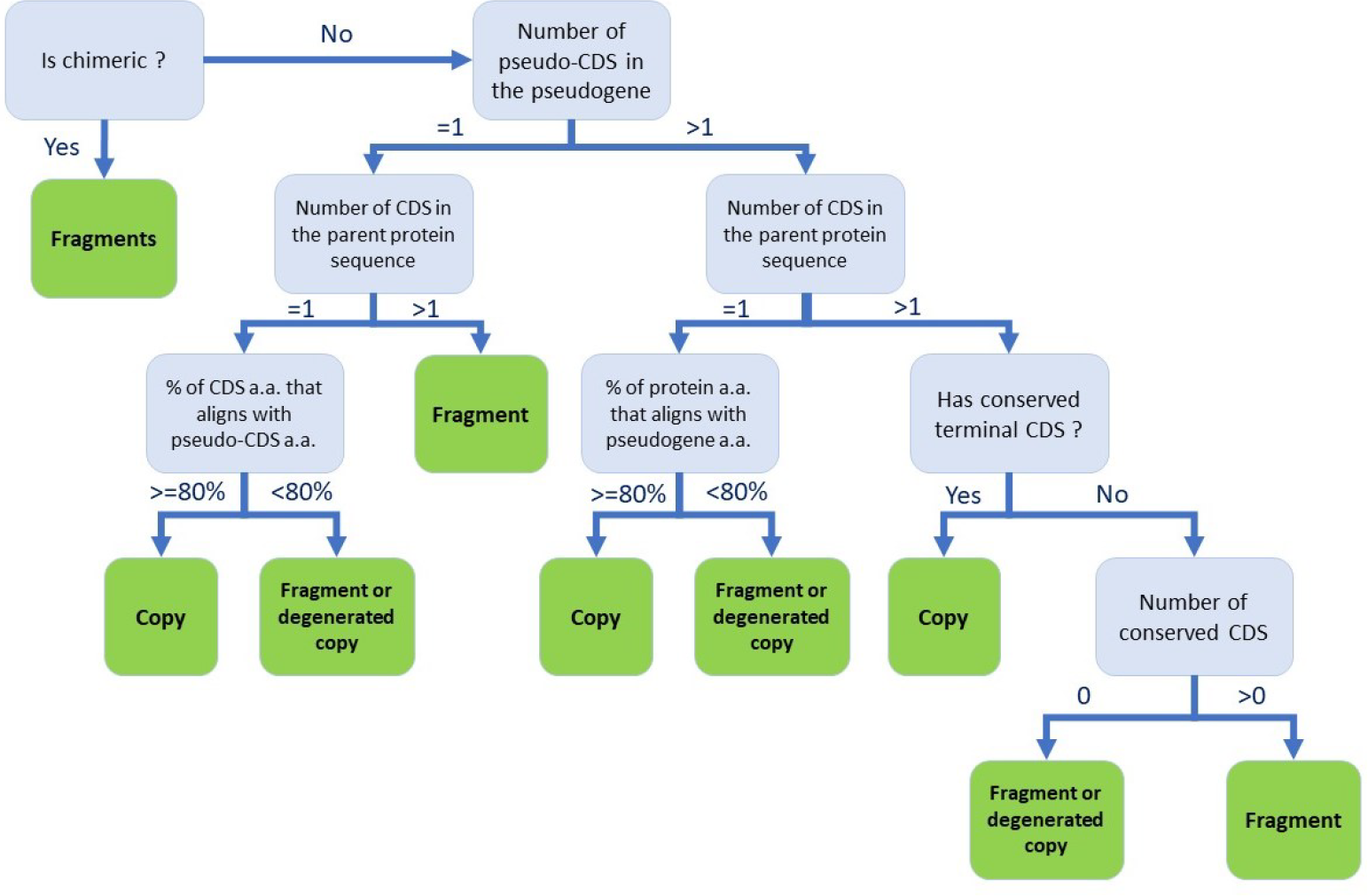
Decision tree to categorize the completeness of pseudogenes predicted by P-GRe. Conserved coding sequence (CDS) are defined as pseudogene (pseudo-)CDS encoding an amino acid sequence producing an alignment containing less than 20% gap with the CDS of a parent protein. These percentages are calculated at the time of alignment corrected by the Lindley-inspired process.

The pseudogene type is determined from intron losses or retentions and the presence or absence of a poly(A) tail. It may be noted “Duplicated pseudogene” if the pseudogene appears to be a duplication of the parent gene, “Retropseudogene” if it appears to be the result of a reverse transcription of the parent gene, or “(Iso)retropseudogene” if it appears to be derived of a reverse transcription of an isoform of the parent gene. This characteristic is also determined using a decision tree (Figure 6). Note that for this step, a poly(A) tail search can be carried out by P-GRe in an area of (maximum) 500 bases following the last position on the 3’ side of a pseudogene. A poly(A) tail is considered present if a series of 20 bases containing at least 15 A is found in that area. Conversely, it is considered absent if no series of 20 bases with more than 7 A is found in this area.

**Figure 6.**
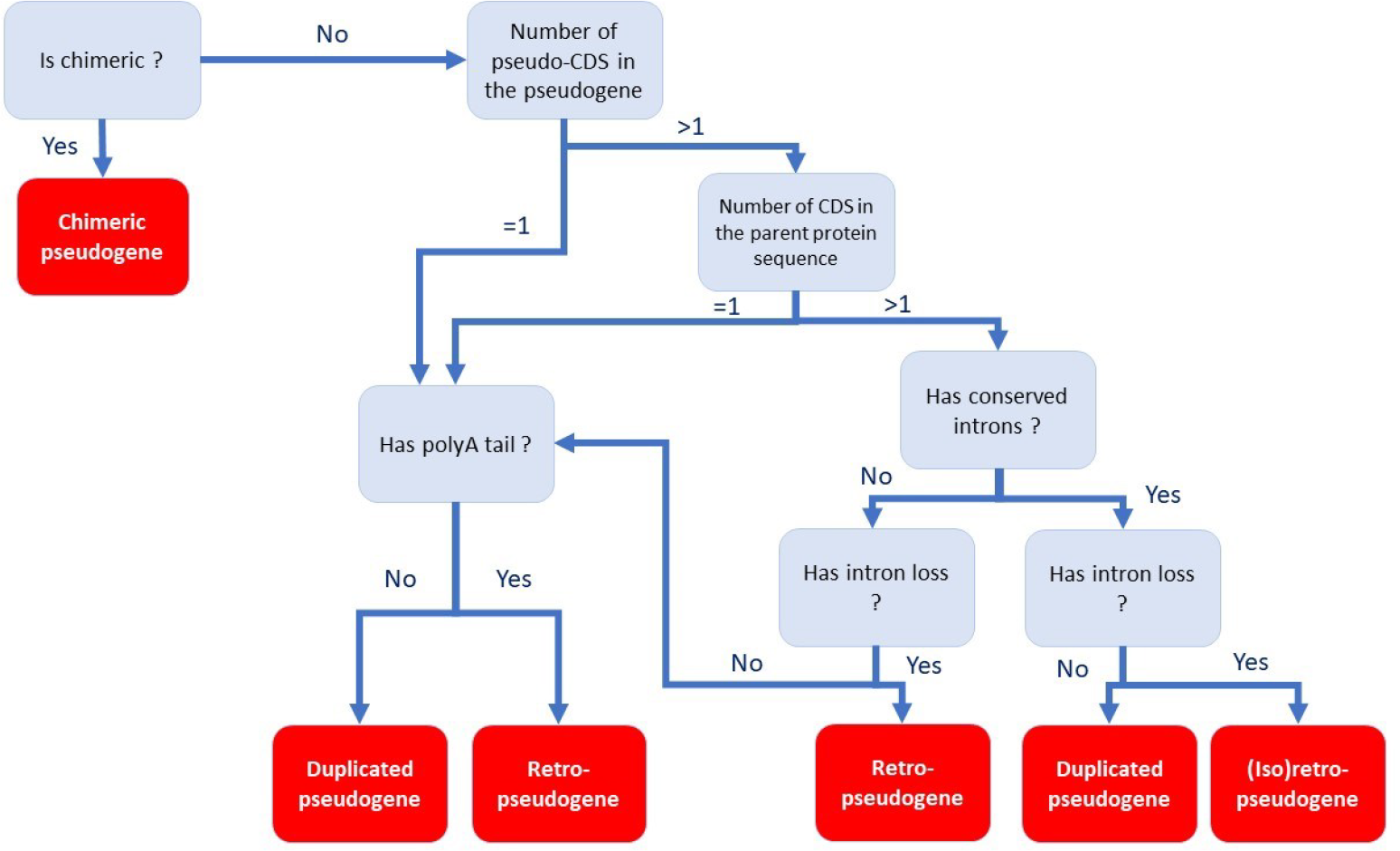
Decision tree to categorize the type of pseudogenes predicted by P-GRe. Intron loss or retention is analyzed at the time of alignments corrected by the Lindley-inspired process.

Furthermore, losses or retention of introns are only sought between two conserved (pseudo-)CDSs, *i*.*e*., CDSs of pseudogenes which produce an alignment with a CDS of a parent protein containing less than 20% gap.

### Quality of predictions and sensibility

The sensitivity of P-GRe was tested using annotation, proteome and genome from *A. thaliana* available on the Ensembl database (TAIR10). Among the 924 known *A. thaliana* pseudogenes, those that overlapped pseudogenes predicted by P-GRe over at least 60% of their length were considered found. Note that this sensitivity metric and the data used for prediction are similar to those used by Xiao *et al*. (Xiao *et al*., 2016), which achieved 78.9% and 81.3% sensitivity with Shiu’s pipeline and PseudoPipe respectively. A similar method was used to identify the predicted pseudogenes that corresponded to transposable elements (TEs, annotated “transposable_element_gene” or “transposable_element” on the TAIR10 annotation). In addition, the sequences corresponding to the elements annotated as “transposable_element_gene” were translated and the sequences virtually encoded by the pseudogenes predicted by P-GRe were locally aligned with these TE sequences using the blastp algorithm.

In addition to sensitivity, the quality of the predictions was also evaluated. All pseudogenes found were aligned locally against the entire proteome of *A. thaliana* using the blastp algorithm, keeping for each pseudogene only the best alignment. From these local alignments, percentage of identity, hit coverage, and total coverage (from the start of the first hit to the end of the last hit) were retained. Each protein sequence that lead to a best alignment was then semi-globally aligned with their matching pseudogene protein sequence using the pairwise2 module, with gap opening and gap extension penalties of 0.5 and 0.1, respectively, and with the BLOSUM62 substitution matrix. From these semi-global alignments, the alignment scores were retained. In the rare cases where a pseudogene predicted by P-GRe overlapped several annotated pseudogenes, the score for each of the TAIR10-annotated pseudogenes was set to the sum of their scores. The same kind of semi-global alignment was used to compare pseudogenes predicted by P-GRe that overlapped known pseudogenes with those that did not overlap and were not associated with TEs.

Note that the prediction quality measurement, described in the previous paragraph, uses both local and semi-global alignments because it is common for pseudogenes to be a ends-truncated copy of a parent gene, or to be highly degenerated, where only a few short preserved and scattered regions remain.

RESULTS & DISCUSSION

### Comparison with known pseudogenes

779 (84.31%) of the 924 annotated pseudogenes in *A. thaliana* were predicted by P-GRe, for a total of 8459 predicted pseudogenes. The quality of the predictions of the 779 pseudogenes predicted by P-GRe was measured and compared to the corresponding annotated pseudogenes. Overall, P-GRe pseudogenes achieved better alignments with their parent proteins than annotated pseudogenes, which is particularly marked by an average alignment score higher by 159.31 points. In concrete terms, over 70% of the pseudogenes predicted by P-GRe match one of the *A. thaliana* proteins with an alignment score higher than that of their known counterparts annotated in TAIR10 (Table 1).

**Table 1.** Measurements of the quality of predictions made by P-GRe and comparison with TAIR10 annotated pseudogenes. The quality of the amino acid sequences virtually encoded by the set of pseudogenes common to P-GRe and the TARI10 annotation was measured by aligning each sequence locally against the entire *A. thaliana* proteome. The identity, coverage rate of the subject protein by hits and total coverage rate (from the first hit to the last) were measured. A semi-global alignment between the query and subject sequences was then performed and the alignment score was recorded.

|  | Identity | Average hit coverage | Average total coverage | Average semi-global alignment score |
| --- | --- | --- | --- | --- |
| P-GRe | 0.64 | 0.56 | 0.81 | 1018.48 |
| TAIR10 | 0.65 | 0.42 | 0.71 | 859.17 |

### TE-related pseudogenes

Of the pseudogenes predicted by P-GRe, 4505 aligned locally with the sequences of known and annotated TEs in TAIR10. Among these TE-associated predictions, the positions of 2186 of them overlap the positions of an annotated transposable element over at least 60% of its length. With regard to the remaining 2319 predicted pseudogenes, it should be noted that their local alignments with the TE genes obtain a low E-value of 7.73e-05 in average. For comparison, pseudogenes that overlap annotated TEs produce alignments with a slightly higher average E-value of 8.77e-05 (Table 2). We therefore assume that, rather than being false positives, a large proportion of these pseudogenes predicted by P-GRe could correspond to still unknown TE genes or TE pseudogenes.

**Table 2.** Comparison of local alignments used to generate pseudogenes similar to transposable elements (TEs). Predicted pseudogenes were considered to overlap a known TE if their positions overlapped at least 60% of a known TE. The right column contains data for predicted pseudogenes that did not overlap a known TE but still produced local alignments with annotated TE sequences.

|  | Predicted pseudogenes overlapping known TE | TE-similar predicted pseudogenes not overlapping any known TE |
| --- | --- | --- |
| Hits average identity | 0.62 | 0.63 |
| Hits average length | 49.44 | 53.91 |
| Hits average E-value | 8.77e-05 | 7.73e-05 |

### Non-TE-related pseudogenes

Among the pseudogenes predicted by P-GRe, 2913 do not cover any genes from TAIR10 annotation, or any transposable elements or pseudogenes, nor do these pseudogenes produce any significant alignment with known transposable elements in *A. thaliana*. These pseudogenes were constructed by P-GRe with hits having an average E-value of 4.94e-04 and an average identity of 64%, which is similar to those corresponding to annotated pseudogenes, as they were constructed from hits with an average E-value of 1.05e-04 and an average identity of 63%. The only notable difference between these two situations is the average length of the hits that enabled them to be generated. Indeed, pseudogenes predicted by P-GRe and not corresponding to any known pseudogene were generated from hits with an average length of 47.77 amino acids, whereas those corresponding to known pseudogenes were generated from hits with an average length of 78.65 amino acids. As a consequence, the average sizes of the complete pseudo-proteins are 100 amino acids and 454 amino acids respectively for previously unknown and known pseudogenes. Nevertheless, when aligning these pseudo-proteins to their respective parent proteins, an average relative alignment score of 3.76 per amino acid is reached, while it is 3.70 per amino acid for the annotated ones (Table 3). Overall, this suggests that P-GRe may be able to detect short pseudogene fragments that were missed in current annotation.

**Table 3.**
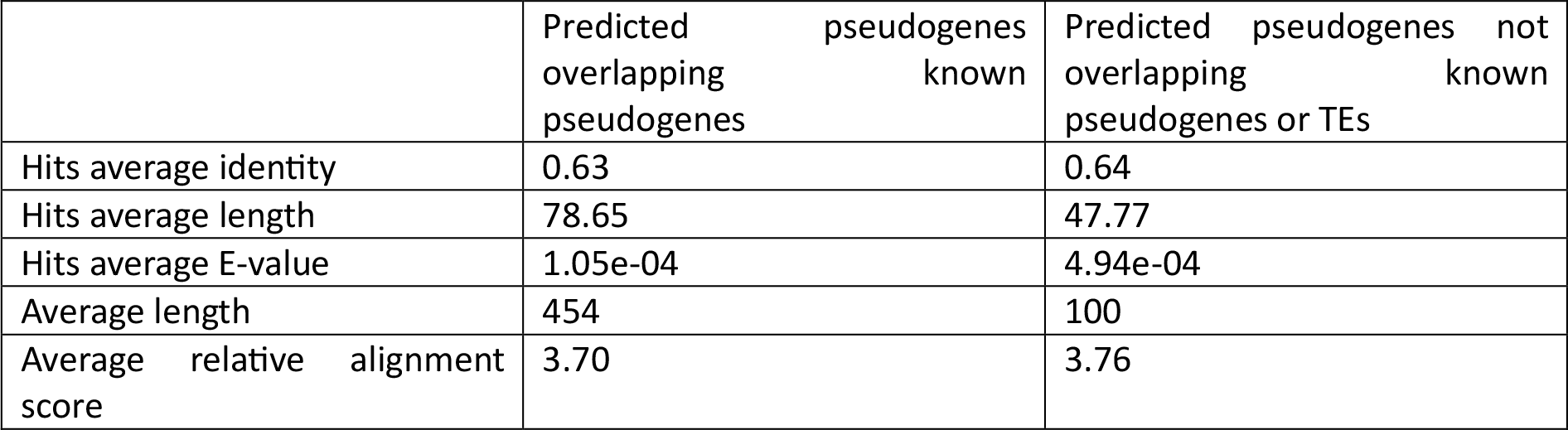
Comparison between pseudogenes predicted by P-GRe corresponding to annotated pseudogenes and the other non-TE-related pseudogenes. The first three lines correspond to average values concerning the local alignments used for generating the pseudogenes. The average length is that of the peptides and proteins virtually encoded by the two categories of pseudogenes, and the average relative alignment score is the average score of the pseudogenes obtained by dividing the score of the semi-global alignment between a pseudogene and its parent gene by the length of the pseudogene.

### Pseudogene type and sequence conservation

The classification of pseudogenes carried out by P-GRe was compared between the three categories of pseudogenes, *i*.*e*. predicted pseudogenes which overlap known pseudogenes, predicted pseudogenes which overlap known TEs, and predicted pseudogenes which overlap neither and whose sequences do not align with the sequences of the known TEs. Striking differences were found between these categories (Table 4). For pseudogenes overlapping known pseudogenes, 19% were noted as full copies, compared to 9% and 7%, respectively, for the other two pseudogene categories. 13% of these pseudogenes were also noted as coming from duplication event, compared to 5% and 4% for the other pseudogene categories. These data seem to demonstrate that the annotations of the pseudogenes of *A. thaliana* are mainly focused on pseudogenes with relatively conserved sequences, close to their parent gene sequences, which could explain the large number of new predicted pseudogenes. In contrast, predicted pseudogenes that were not annotated appeared more heavily degraded, with 85% of them classified as fragments or degraded copies, compared to 51% and 40% for pseudogenes overlapping annotated pseudogenes or annotated TEs, respectively. In addition, the type of 80% of them could not be determined, compared to 48% and 37% for the pseudogenes overlapping known TEs and those not overlapping any known TEs or annotated pseudogenes, respectively.

**Table 4.** Type and completeness of the different categories of predicted pseudogenes.

|  | Predicted pseudogenes overlapping known pseudogenes | Predicted pseudogenes overlapping known TEs | Predicted pseudogenes not overlapping known pseudogenes or TEs |
| --- | --- | --- | --- |
| Completeness |  |  |  |
| Copy | 19% | 9% | 7% |
| Fragment | 35% | 23% | 56% |
| Chimeric fragments | 29% | 50% | 8% |
| Fragment or highly degenerated copy | 16% | 17% | 29% |
| Type |  |  |  |
| Chimeric pseudogene | 29% | 50% | 8% |
| Duplicated pseudogene | 13% | 5% | 4% |
| (Iso)retropseudogene | 3% | 1% | 0% |
| Retropseudogene | 7% | 6% | 7% |
| Undetermined | 48% | 37% | 80% |

Interestingly, the proportion of pseudogene noted as chimeric was much higher in predicted pseudogenes overlapping TEs (50%), than in pseudogenes overlapping annotated pseudogenes (29%) or in non-annotated pseudogenes (8%). This result was expected, as TEs have a role in the formation of this type of pseudogene. However, it should be noted that the number of pseudogenes noted as chimeric may be exaggerated due to pseudogenes having diverged so much from their parent that they have similarities with several different genes. Chimeric pseudogenes aside, we also found that 10% of retropseudogenes originated from a retrotransposition event on an alternative mRNA (iso-retrotransposon).

## Acknowledgements

The authors are thankful to the Paul Sabatier-Toulouse 3 University and to the Centre National de la Recherche Scientifique (CNRS) for granting their work. SC is the recipient of a fellowship from the “École Universitaire de Recherche (EUR)” TULIP-GS (ANR-18-EURE-0019). This study is set within the framework of the “Laboratoires d’Excellences (LABEX)” TULIP (ANR-10-LABX-41).

